# Thermophilic PHB depolymerase of *Stenotrophomonas* sp., an isolate from the plastic contaminated site is best purified on Octyl-Sepharose CL-4B

**DOI:** 10.1101/618967

**Authors:** R Z Sayyed, S J Wani, S S Shaikh, Helal F. Al-Harthi, Asad Syed, Hesham Ali El-Enshasy

**Affiliations:** Department of Microbiology, PSGVP Mandal’s, Arts, Science, and Commerce College, SHAHADA, Maharashtra 425409 India; Department of Botany and Microbiology, College of Science, King Saud University, Riyadh, 11451, Saudi Arabia; School of Chemical and Energy Engineering, Faculty of Engineering, Universiti Teknologi Malaysia (UTM), Skudai, Johor Bahru, Malaysia; City of Scientific Research and Technology Applications, New Burg Al Arab, Alexandria, Egypt

**Author notes:** Corresponding author R.Z. Sayyed. These authors contributed equally to this work (S J Wani, S S Shaikh). These authors contributed equally to this work (Helal F. Al-Harthi, Asad Syed, Hesham Ali El-Enshasy).

**Keywords:** Biodegradable polymer, PHB depolymerase, Thermophilic, *Stenotrophomonas* sp

## Abstract

There are numerous reports on PHB depolymerases produced by a wide variety of microorganisms isolated from various habitats, however, reports on PHB depolymerase isolated from plastic contaminated sites are scares. Thermophilic PHB polymerase produced by isolates obtained from plastic contaminated sites is expected to have better relevance for its application in plastic/ bioplastic degradation. Although PHB has attracted commercial significance, the inefficient production and recovery methods, inefficient purification of PHB depolymerase and lack of ample knowledge on PHB degradation by PHB depolymerase have hampered its large scale commercialization. Therefore, to ensure the biodegradability of biopolymers, it becomes imperative to study the purification of the biodegrading enzyme system. We report the production, purification, and characterization of extracellular PHB depolymerase from *Stenotrophomonas* sp. RZS 7 isolated from a plastic contaminated site. The isolate produced extracellular poly-β-hydroxybutyrate (PHB) depolymerase in the mineral salt medium at 30oC during 4 days of incubation under shake flask condition. Purification of the enzyme was carried out by three different methods using PHB as a substrate. Purification of PHB depolymerase by ammonium salt precipitation, column chromatography, and solvent purification method was successfully carried out. Among the purification method tested, the enzyme was best purified by column chromatography on Octyl-Sepharose CL-4B column with maximum (0.7993 U/mg/ml) purification yield. The molecular weight of purified PHB depolymerase (40 kDa) closely resembled with PHB depolymerase of *Aureobacterium saperdae*.

## Introduction

Poly-β-hydroxy alkanoates (PHAs) or Poly-β-hydroxybutyrate (PHB) are accumulated as a source of food and energy by a wide variety of bacteria growing under nitrogen-deficient but conditions and is mobilized during nutrient stress under the influence of PHB depolymerase (1-6). PHA/PHB is considered as the best eco-friendly and renewable alternative to the synthetics petrochemical plastics because of its similar properties to synthetic plastic (7-8) besides being thermoplastic and biodegradable in nature. Because of such useful properties, it has attracted commercial interest for use as the best alternative to the hazardous synthetic petrochemical polymers and hence it has been successfully commercialized. During last decades much research has been devoted towards distribution and occurrence of PHB degraders and studies on different PHB depolymerases. Jendrossek and Handrick (9) reported that PHB depolymerases are responsible for extracellular PHB degradation. Extracellular PHB depolymerases of *Aspergillus fumigatus* Pdf1 (10), *A. Saperdae* (11), *Thermus thermophiles* HB8 (12), *Streptomyces bangladeshensis* 77T-4 (13), *Penicillium simplicissimum* LAR13 (14), *Acidovorax* sp. TP4 (15), *Emericellopsis minima* W2 (16) has been isolated and purified.

However, the organism isolated from the plastic contaminated site and having the ability to degrade PHB may be a potential source of dynamic PHB depolymerase. However, inefficient production and recovery process of PHB, inefficient purification of PHB depolymerase and lack of ample knowledge on PHB degradation by PHB depolymerase have hampered the large scale commercialization of PHB. Therefore, to ensure the biodegradability of biopolymers, it becomes imperative to study the purification of biodegrading enzyme system (16).

The present paper reports the production, purification, and characterization of extracellular PHB depolymerase of *Stenotrophomonas* sp. RZS 7 isolated from a plastic contaminated site.

## Material and Methods

### PHB

All the experiment was carried out using PHB powder. PHB was obtained from Sigma-Aldrich, Germany.

### Source of bacterium

*Stenotrophomonas* sp. RZS 7 isolated from plastic contaminated site was previously identified (17) and used in the present study as a source of PHB depolymerase.

### Production of PHB depolymerase

Production of PHB depolymerase was carried out at shake flask level by growing *Stenotrophomonas* sp. RZS 7 in minimal medium (MM) containing PHB, 0.15%; K_2_HPO_4_, 0.7 g; KH_2_PO_4_, 0.7 g; MgSO_4_, 0.7 g; NH_4_Cl, 1.0 g; NaNO_3_, 1.0 g; NaCl, 5 mg; FeSO_4_, 2 mg, ZnSO_4_, 7 mg in 1 L of distilled water (14) at 120 rpm for 4 days at 30^°^C.

### PHB depolymerase assay

Following 4 day’s incubation at 30^°^C and 120 rpm, MM broth was centrifuged a 10,000 rpm for 15 min and the supernatant was assayed for PHB depolymerase (12). For this purpose PHB granule (substrate for PHB depolymerase) were sonicated (20 kHz for 15 min) and suspended in 50 mM Tris-HCl buffer (pH 7.0) and 150 μg/ml of this suspension and 2 mM CaCl_2_ was added in 50mM Tris-HCl buffer (pH 7.0) followed by the addition of culture supernatant (0.5 ml). Enzyme activity was spectrophotometrically measured at 650 nm as a decrease in the PHB turbidity. One unit of PHB depolymerase was defined as the quantity of enzyme required to decrease the absorbance by 0.1 /min.

### Purification of PHB depolymerase

Having confirmed the presence of PHB depolymerase in cell-free supernatant of MM broth, the broth was subjected for purification of the enzyme by three approaches as follows

### Ammonium salt precipitation

The crude enzyme in the culture supernatant was precipitated by adding solid ammonium sulfate with continuous stirring at 4^°^C for 1 h. The precipitate was dissolved in the Tris-HCl buffer (pH 7) supernatant and dissolved precipitate was transferred in a separate dialysis bag and allowed for overnight dialysis in chilled phosphate buffer (13). The protein concentration of dialyzed supernatant, as well as dialyzed precipitate, was measured by the Lowry method (18). PHB depolymerase activity of dialyzed supernatant and dialyzed precipitate was measured as described earlier.

### Solvent purification method

The culture supernatant of the isolate was centrifuged at 10,000 rpm for 20 min. The residue obtained was dissolved in pre-chilled 1:1 acetone ethanol mixture, shaken well and kept in a water bath at 50^°^C until all solvent is evaporated. The pellet obtained after evaporation was dissolved in Tris-HCl buffer (pH 7). PHB depolymerase activity of pellet, as well as supernatant, was carried out as described.

### Column chromatography

The cell-free supernatant of the isolate was applied onto an Octyl-Sepharose CL-4B column pre-equilibrated with 50 mM glycine NaOH buffer (pH 9.0) and eluted with a gradient of 0 to 50% ethanol (12), the fractions were collected and PHB depolymerase activity of each fraction was determined as described.

### Molecular weight determination of purified PHB depolymerase

To measure the molecular weight of purified PHB depolymerase, SDS PAGE (Sodium Dodecyl Sulfate Polyacrylamide Gel Electrophoresis) was performed with standard molecular weight markers such as phosphorylase b (82.2 kDa), bovine serum albumin (64.2 kDa), egg albumin (48.8 kDa), carbonic anhydrase (37.1 kDa), trypsin inhibitor (25.9 kDa), lysozyme (19.4 kDa), lysozyme (14.8 kDa) and lysozyme (6.0 kDa). The protein concentration of purified band was measured by the Lowry method with bovine serum albumin (BSA) as a standard (14).

### Statistical analysis

All the experiments were performed in triplicate and the mean of three replicates was considered. Each mean value was subjected to Student’s *t-*test and values of *P* ≤ 0.05 were taken as statistically significant (19).

## Results and Discussion

### Production of PHB depolymerase

After 4 days’ incubation at 30^°^C at 120 rpm in MM, *Stenotrophomonas* sp. RZS 7 yielded 0.721 U/ml PHB depolymerase.

### Purification of PHB depolymerase

#### Ammonium salt precipitation

With increasing concentrations of ammonium sulfate in the cell-free culture supernatant of *Stenotrophomonas* sp. RZS 7, increased precipitation of protein was observed, maximum proteins were precipitated with 70% ammonium sulfate concentration (Table 1). The protein concentration and PHB depolymerase specific activity and enzyme activity of dialyzed precipitate of *Stenotrophomonas* sp. RZS 7 were 0.219 mg/ml, 0.7031 U/mg/ml and 0.154 U/ml respectively. Zhou et al. (20) have reported precipitation of PHB depolymerase of *Escherichia coli*, and *Penicillium* sp. DS9701-D2 by using 70 and 75% of ammonium sulfate Shivkumar et al. (21) have reported efficient precipitation of PHB depolymerase of *Penicillium citrinum* S2 with 80% of ammonium sulfate.

**Table 1 –.** Purification of PHB depolymerase of *Stenotrophomonas* sp. RZS7 on Octyl sepharose column.

| **Fraction** | **Total activity**<br>**(U/ml)** | **Total protein**<br>**(mg/ml)** | **Specific activity**<br>**(U/mg/ml)** | **% activity** |
| --- | --- | --- | --- | --- |
| 1 | 0.027 (0.027) | 0.031 (0.010) | 0.0782 (0.032) | 11.13 (0.012) |
| 2 | 0.101 (0.011) | 0.153 (0.037) | 0.3781 (0.041) | 48.31 (0.039) |
| 3 | 0.247 (0.015) | 0.309 (0.074) | 0.7993 (0.090) | 79.93 (0.092) |
| 4 | 0.017 (0.001) | 0.015 (0.009) | 0.0106 (0.011) | 04.01 (0.012) |
| 5 | 0.003 (0.001) | 0.006 (0.005) | 0.0011 (0.008) | 00.10 (0.002) |

Values were taken to be statistically significant at $P \leq 0.05$
Values are the mean of three replicates

#### Solvent purification method

Solvent purification of PHB depolymerase of *Stenotrophomonas* sp. RZS 7, retained only 51.14 % and 38.81% enzyme activity in pellet and supernatant (Table 2).

**Table 2 –.** Purification profile of PHB depolymerase by various methods.

| Isolate | Purification method | Enzyme activity (U/mg/ml) |
| --- | --- | --- |
| *Stenotrophomonas* sp. RZS 7 | Salt precipitation | 0.7031 (0.090) |
|  | Solvent purification | 0.1120 (0.022) |
|  | Column chromatography | 0.7993 (0.099) |

Values were taken to be statistically significant at $P \leq 0.05$
Values are the mean of three replicates

The protein concentration and PHB depolymerase specific activity and enzyme activity in pellet and supernatant were 0.219 mg/ml, 0.5114 U/mg/ml and 0.3881 U/ml respectively. This significant loss of enzyme activity may be due to precipitation resulting in denaturation of proteins by solvent system. Thus the solvent purification method proved insignificant vis-à-vis other purification methods.

#### Column chromatography

The dialyzed precipitate of *Stenotrophomonas* sp. RZS 7 obtained after ammonium salt precipitation when applied onto an Octyl-Sepharose CL-4B column pre-equilibrated with 50 mM glycine NaOH buffer (pH 9.0) and eluted with ethanol, yielded 5 fractions. Among all the fractions analyzed for PHB depolymerase enzyme activity, 3^rd^ fractions showed maximum enzyme activity (Table 1). PHBV depolymerase of *Bacillus* sp. AF3 and *Streptoverticillium kashmirense* AF1 have also been purified on Sephadex G-75 (22). Kim et al. (23) have also purified PHB depolymerase of *Emericellopsis minima* W2 and *Streptomyces* sp. KJ-72 on Sephadex G-100 and Sephadex G-150 respectively. Papaneophytou et al. (12) and Hsu et al. (13) have isolated and purified extracellular PHB depolymerases of *Thermus thermophiles* HB8 and *Streptomyces bangladeshensis* 77T-4 respectively using column chromatography.

Among all three purification methods, purification on Octyl sepharose CL-4B column gave good purification yield (Table 1), more enzyme activity and more specific activity. The protein concentration and PHB depolymerase specific activity and enzyme activity were 0.247 mg/ml, 0.7993 U/mg/ml and 0.309 U/ml respectively.

### Molecular weight determination of purified PHB depolymerase

The protein fraction of *Stenotrophomonas* sp. RZS 7 obtained from column chromatography having maximum PHB depolymerase enzyme activity showed single protein band in SDS PAGE having a molecular mass of approximately 40 kDa lying between molecular weight markers of 37.1 and 48.8 (Fig 1). Sadocco et al. (11) have also reported the molecular mass of PHB depolymerase from *Aureobacterium saperdae* ranges from 42.7 kDa analyzed by SDS PAGE.

## Conclusion

The present attempt provided PHB degrading *Stenotrophomonas* sp. RZS 7 which utilized PHB as a sole source of carbon under the influence of extracellular PHB depolymerase. The enzyme acted as a key enzyme responsible for biodegradation of PHB. Purification of PHB depolymerase by solvent purification resulted in protein precipitation and denaturation of the enzyme. Column chromatography by using octyl-Sepharose CL-4B came out as efficient and best purification method as it purification yielded maximum protein, maximum enzyme activity, and maximum specific activity. The molecular weight of purified PHB depolymerase of *Stenotrophomonas* sp. RZS 7 (40kDa) matched with a molecular weight of *Aureobacterium saperdae*.

## Competing interests

All authors declare no conflict of interest.

## Author Contributions

### Conceptualization and drafting

R. Z. Sayyed

### Investigation

S. J. Wani and S S Shaikh

### Data curation

Abdullah A. Alyousef, Abdulaziz Alqasim

### Funding acquisition

Abdullah A. Alyousef, Abdulaziz Alqasim, Hesham Ali El-Enshasy

### Data Availability Statement

All relevant data are within the paper.

## Funding

This research is supported by grant No. RG-1440-053 by the Deanship of Scientific Research at King Saud University, Saudia Arabia and the support of MOE and UTM-RMC (Malaysia) through HICOE grant No. R.J130000.7846.4J262.

